# Ecological differences among hydrothermal vent symbioses may drive contrasting patterns of symbiont population differentiation

**DOI:** 10.1101/2022.08.30.505939

**Authors:** Corinna Breusing, Yao Xiao, Shelbi L. Russell, Russell B. Corbett-Detig, Sixuan Li, Jin Sun, Chong Chen, Yi Lan, Pei-Yuan Qian, Roxanne A. Beinart

## Abstract

The intra-host composition of horizontally transmitted microbial symbionts can vary across host populations due to interactive effects of host genetics, environmental and geographic factors. While adaptation to local habitat conditions can drive geographic subdivision of symbiont strains, it is unknown whether these patterns are universal and how differences in ecological characteristics among host-symbiont associations influence the genomic structure of symbiont populations. To address this question, we sequenced metagenomes of different populations of the deep-sea mussel *Bathymodiolus septemdierum*, which are common at Western Pacific deep-sea hydrothermal vents and show characteristic patterns of niche partitioning with sympatric gastropod symbioses. *Bathymodiolus septemdierum* lives in close symbiotic relationship with sulfur-oxidizing chemosynthetic bacteria but supplements its symbiotrophic diet through filter-feeding, enabling it to occupy ecological niches with less exposure to geochemical reductants. Our analyses indicate that symbiont populations associated with *B. septemdierum* show structuring by geographic location, but that the dominant symbiont strain is uncorrelated with vent site. These patterns are in contrast to co-occurring *Alviniconcha* and *Ifremeria* symbioses that exhibit greater symbiont nutritional dependence and occupy habitats with higher spatial variability in environmental conditions. Our results suggest that relative habitat homogeneity combined with sufficient symbiont dispersal and genomic mixing might promote persistence of similar symbiont strains across geographic locations, while mixotrophy might decrease selective pressures on the host to affiliate with locally adapted symbiont strains. Overall, these data contribute to our understanding of the potential mechanisms influencing symbiont population structure across a spectrum of marine microbial symbioses that vary in ecological niche and relative host dependency.

## Introduction

In the ocean, mutualistic symbioses between animals and bacteria are predominantly established through symbiont acquisition from the environment [1]. The composition of environmentally acquired symbionts within and across host populations is typically influenced by a combination of host genetic makeup, environmental selection, physical barriers to symbiont dispersal, symbiont competition and horizontal gene transfer, including homologous recombination (i.e., the exchange of homologous genetic material between bacterial strains or species) [2–4]. For instance, while host specificity, ecological partitioning or dispersal limitations may increase genetic divergence among symbiont populations, frequent recombination may promote genetic cohesiveness within a symbiont species. However, in most cases, the relative importance of these factors in shaping the genetic structure of symbiont populations remains unclear. Similarly, it is unknown how differences in ecological characteristics among symbiotic partnerships may impact patterns of symbiont population subdivision. Further work is necessary to untangle the effects of neutral and adaptive processes, as well as the dynamics of host-symbiont interaction, on horizontally transmitted symbiont populations.

Hydrothermal vents are dominated by invertebrates that obligately rely on the chemosynthesis of their bacterial symbionts for nutrition in the otherwise food-limited deep sea. Most symbiotic vent animals transmit their symbionts horizontally [5], such that at some early developmental stage, host animals must acquire their bacterial symbiont from the environment to survive. In these systems, where aposymbiotic larvae disperse between geographically distant and environmentally variable island-like vent fields, the acquisition of a local symbiont strain after settlement is thought to be common [6, 7]. Indeed, studies show that vent species typically host distinct symbiont strains that vary by geographic site [7–13]. Our previous work on *Alviniconcha* and *Ifremeria* symbioses from the Lau Basin indicated that site-specific variation in symbiont populations is driven by environmental selection [7], though it is currently unknown whether these processes apply to other hydrothermal vent symbioses that differ in their ecological characteristics.

Owing to their ecological and evolutionary success, bathymodioline mussels belong to the dominant fauna at deep-sea hydrothermal vents worldwide [14], with the genus *Bathymodiolus* representing the most prominent member of the group. In the majority of cases, *Bathymodiolus* species host thiotrophic gammaproteobacterial symbionts that use reduced sulfur compounds as energy sources for chemosynthesis [15]. *Bathymodiolus septemdierum* is widespread in the Indo-Pacific Ocean and although various names have been proposed for different morphotypes, including *B. septemdierum*, *B. brevior*, and *B. marisindicus*, both morphometric and genetic data have revealed that they represent populations of the same species, leading to the other names being synonymized with *B. septemdierum* [16–18]. In the Western Pacific, this mussel forms expansive beds that are typically arranged in concentric rings around patches of sympatric gastropods of the genera *Alviniconcha* and *Ifremeria* [19]. Fine-scale habitat partitioning among these holobionts is likely driven by contrasting thermal tolerances and sulfide requirements for chemosynthesis, resulting in differential exposure of hosts and symbionts to available geochemical reductants: *Alviniconcha* holobionts usually encounter the highest and most variable sulfur concentrations, while *Bathymodiolus* holobionts experience the lowest and most stable levels of sulfur compounds [19, 20]. The ability of *Bathymodiolus* symbioses to thrive under diminished sulfide availability is hypothesized to be partly related to the hosts’ mixotrophic feeding mode [19, 21], which would decrease dependence on optimal chemosynthetic production rates. Indeed, stable isotope and gill structure analyses suggest that *B. septemdierum* is unlikely to fulfill its dietary requirements solely by assimilating symbiont-derived organic carbon, especially given that symbiont densities appear to be too low to provide the host with sufficient nutrition [22].

In this study we sequenced metagenomes of *B. septemdierum* populations from four different vent sites in the Lau Basin to assess the population genetic structure and strain-level functional variation of its thiotrophic symbiont, *Candidatus* Thiodubiliella endoseptemdiera [4], in relation to host genotype, environmental factors and geographic location. To examine if ecological differences and homologous recombination might play a role in symbiont geographic partitioning, we subsequently compared the results to patterns in co-occurring *Alviniconcha* and *Ifremeria* symbioses [7] that show strong contrasts in their niche preferences.

## Materials and Methods

### Sample collection, DNA extraction and sequencing

Mussel samples were collected from the Tu’i Malila (n = 40), ABE (n = 34), Tahi Moana (n = 11) and Tow Cam (n = 20) vent sites with the remotely operated vehicles (ROV) *ROPOS* and *Jason* during research cruises to the Eastern Lau Spreading Center (ELSC), Southwest Pacific, in 2009 and 2016 (Fig. 1A; Table S1). The sites sampled are representative of the range of geochemical habitat conditions *B. septemdierum* holobionts experience within the Lau Basin. Upon ROV recovery, symbiont-bearing gill tissue of each individual was dissected on board ship and frozen or stored in RNALater™ (Thermo Fisher Scientific, Inc., Waltham, MA, USA) at – 80°C until further analysis. DNA of 80 specimens was extracted from gill tissue snips with the E.Z.N.A Mollusc DNA Kit (Omega Bio-tek, Norcross, GA) at the University of Rhode Island and sent for plexWell384 library preparation and NovaSeq6000 2x150 bp sequencing at Psomagen, Inc. (Rockville, MD, USA) (Table S1). A set of an additional 25 specimens was extracted with the DNeasy Blood & Tissue Kit (Qiagen, Hilden, Germany) at the Hong Kong University of Science and Technology and sent for metagenomic library preparation and NovaSeq6000 2x150 bp sequencing at Novogene, Co. (Beijing, China) (Table S1), resulting in a total of 105 samples for analysis. PERMANOVAs suggested that methodological differences had negligible effects on patterns of symbiont genetic variation relative to biological factors (6.14% *versus* 23.72 % explained variation), confirming the validity of combining the two metagenomic datasets (Table S2). Raw reads for each individual were adaptor-clipped and quality-trimmed with TRIMMOMATIC [23] before analysis. Host species identities were determined morphologically and through assembly of the mitochondrial genome of each sample (see below).

**Fig. 1.**
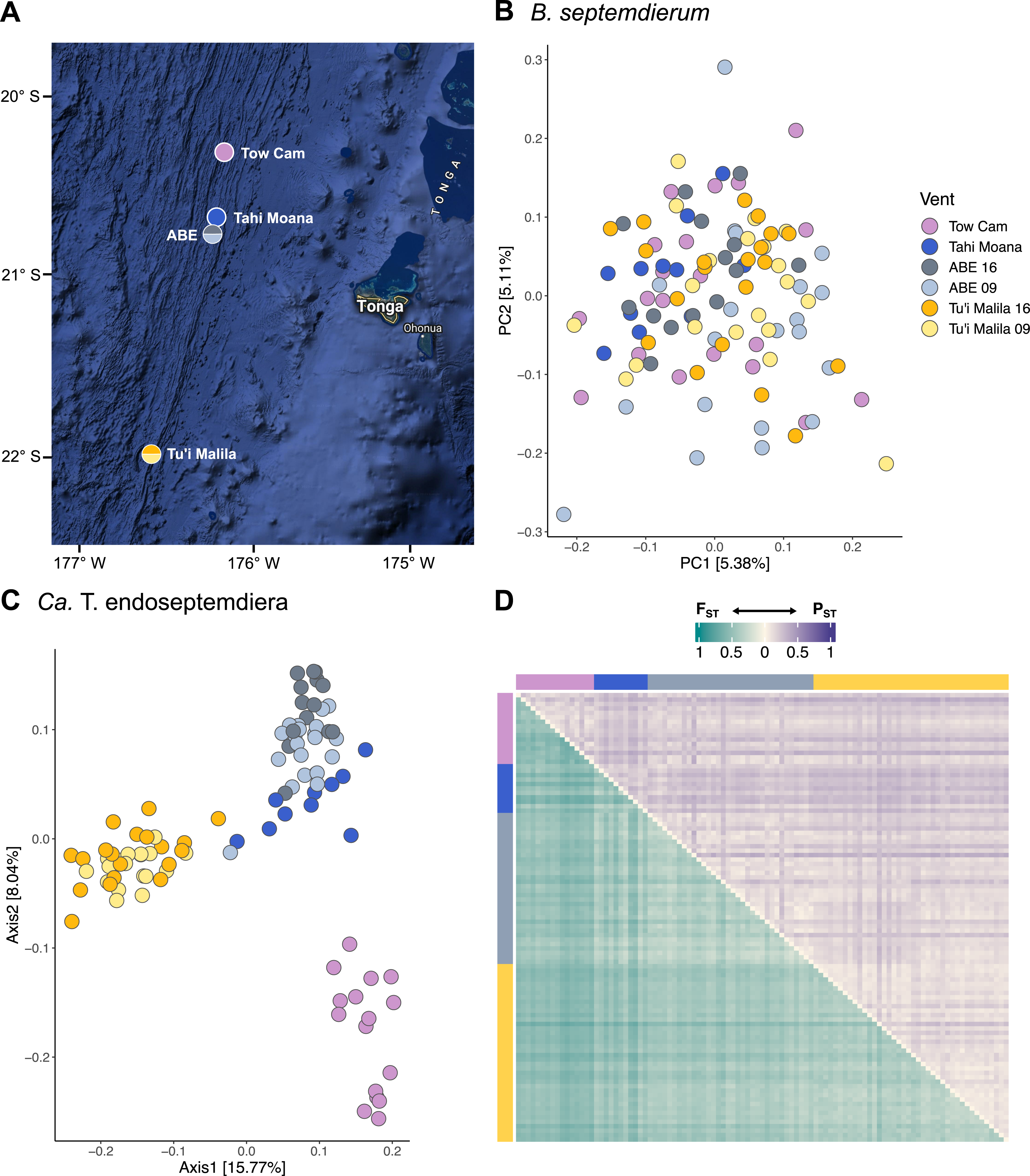
Sampling locations of *Bathymodiolus septemdierum* in the Eastern Lau Spreading Center (A) and genetic structure of host (B) and symbiont (C) populations across vent sites. Vent locations are between 9 and 191 km apart. Host populations are genetically undifferentiated, while moderate structuring by geography is observed in the symbiont. (D) Pairwise F_ST_ values (lower diagonal) and P_ST_ values (upper diagonal) among symbiont populations of single mussel hosts. Compared to patterns observed on a geographic level, differentiation on a per-sample level is notably higher, indicating that symbiont genetic diversity is strongly driven by variation among host individuals. Map data ©2023 Google.

### Host transcriptome generation and population genomic analyses

To determine host population genomic structure, we *de novo* assembled the gill transcriptome of *B. septemdierum* and used the resulting transcripts as reference for variant identification. Total RNA of one RNALater™-preserved sample each from Tu’i Malila, ABE and Tahi Moana was extracted with the Zymo Direct-zol RNA Miniprep Kit (Zymo Research, Inc., Irvine, CA, USA). PolyA-enriched, 150-bp paired-end RNA-seq libraries were prepared with the TruSeq Stranded mRNA Kit (Illumina, Inc., San Diego, CA, USA) and sequenced on a NovaSeq 6000 instrument to a depth of 100 M reads per library by Psomagen, Inc. (Rockville, MD, USA). After initial quality checks with FASTQC (https://www.bioinformatics.babraham.ac.uk/projects/fastqc/), raw reads from all three samples were trimmed with TRIMMOMATIC, error-corrected with RCORRECTOR [24], filtered with TRANSCRIPTOMEASSEMBLYTOOLS (https://github.com/harvardinformatics/TranscriptomeAssemblyTools), and co-assembled with TRINITY in PASAFLY mode [25]. Transcript annotation and further curation, including removal of transcript redundancies, followed the procedure in Breusing et al. [7]. Transcriptome completeness was assessed with BUSCO [26] based on mollusk-specific conserved orthologous genes. Single nucleotide polymorphisms (SNPs) among the 105 sequenced individuals were subsequently estimated by mapping the metagenomic reads against the reference transcriptome and assessing genotype likelihoods with ANGSD [27] based on a model correcting for deviations from Hardy-Weinberg equilibrium (see ref. [7] for details). To evaluate host population genomic differentiation between vent sites we determined pairwise fixation indices (F_ST_s) using the method described in Bhatia et al. [28] and performed principal component analyses of host covariance matrices with the STATS and GGPLOT2 packages in R [29, 30].

### Symbiont pangenome reconstruction and population genomic analyses

To assess symbiont population genomic structure, we reconstructed a pangenome for *Ca.* Thiodubiliella endoseptemdiera out of individual metagenome-assembled genomes (MAGs) that we used as scaffold for read mapping and variant calling. Using a pangenome instead of a single reference genome allowed us to address limitations of individual assemblies, such as the risk of missing accessory genes. Filtered reads from metagenomic sequencing were assembled with METASPADES [31] using kmers from 21 to 121. Binning of MAGs was done following the METAWRAP [32] pipeline and other binning procedures outlined in Breusing et al. [7]. MAGs were taxonomically assigned with GTDB-TK [33] and evaluated for species-level genomic similarity with FASTANI [34]. Ninety-one *Ca.* T. endoseptemdiera MAGs with CHECKM [35] completeness scores > 85% and contamination scores < 10% were considered further for pangenome reconstruction with PANAROO [36]. Genetic variants between symbiont populations were identified by comparing metagenomic reads of all 105 samples against the *Ca.* Thiodubiliella pangenome with FREEBAYES [37] as in Breusing et al. [7]. The identified variants were subsequently used to infer symbiont population structure among vent sites through calculation of pairwise F_ST_s with SCIKIT-ALLEL (https://github.com/cggh/scikit-allel) and principal coordinate analyses with the APE package in R [38]. F_ST_ values between symbiont populations of individual mussel hosts and intra-host nucleotide diversities were calculated after Romero Picazo et al. [39]. Statistical differences between genome-wide intra-host nucleotide diversities among vent sites were determined via pairwise t-tests in R after confirming assumptions of normality. The false discovery rate procedure was applied to correct for multiple testing. To account for differences in variant detection probability within the pangenome, we further calculated the pangenome fixation index (P_ST_) based on gene diversities between vent sites and mussel hosts after Romero Picazo et al. [40]. Associations of symbiont haplotypes with host genotypes, geography and environment were assessed through redundancy analyses and Mantel tests with the VEGAN and NCF packages [41, 42]. Geochemical data for these analyses were taken from ref. [7]. Pairwise gene content differences between symbiont populations from different vent sites were evaluated with PANPHLAN [43] based on coverage profiles of mapped metagenomic reads against the *Ca*. Thiodubiliella pangenome. Genes that were present/absent in ∼90% of samples from a given location were included in further analyses.

### Symbiont strain decomposition and analysis

To quantify the abundance and number of distinct symbiont strains within each mussel host, we inferred strain haplotypes on co-assembly graphs with STRONG [44]. All samples were normalized to the same number of reads before analysis (∼4.1 M reads/sample). STRONG was run by evaluating multiple kmers for assembly (33, 55, 77) and using METABAT2 [45] for binning. MAGs with qualities > 0.85 were considered for strain analysis. The BAYESPATHS algorithm for strain resolution was run in five iterations, using five rounds of gene filtering and allowing up to 15 strains to be detected. For comparative purposes, we also ran DESMAN [46] on the metagenomic data within the STRONG pipeline (parameters: --nb_haplotypes 12 -- nb_repeat 5 --min_cov 1). Phylogenetic relationships among strains were assessed with FASTTREE’s Approximately-Maximum Likelihood method [47]. To evaluate if strain haplotypes represented discrete or cohesive units, we calculated allele frequency spectra (AFS) for each sample from variant calls performed for individual host-associated populations with FREEBAYES, using the same parameters as above except that no threshold on the minimum alternate allele fraction (-F) was applied and minimal post-filtering was performed to prevent bias in the shape of the AFS. Strain abundance heatmaps and phylogenetic trees were produced with the COMPLEXHEATMAP [48] and GGTREE [49] packages in R.

To confirm that strain detection across samples was not markedly influenced by potential sample cross-contamination during processing or sequencing, we evaluated the mitochondrial heterogeneity in each sample. Due to bottleneck effects during vertical transmission, intra-individual mitochondrial diversity should be naturally low, though elevated levels are possible in bivalves in the case of doubly uniparental inheritance where two divergent lineages of female (F) and male (M) mitogenomes are present [50]. Individual host mitochondrial genomes were assembled with MITOBIM [51] using a previously assembled *B. septemdierum* (AP014562) mitogenome as bait for mitochondrial reconstruction. The mitogenome of sample R1931-6_463 could not be resolved through this approach and was reassembled with MITOBIM using the sample’s *COI* sequence as seed. Low frequency variants were then called with LOFREQ [52], using default filters for coverage and strand bias, a minimum mapping quality of 30 and a minimum base quality of 20.

### Homologous recombination among symbiont genomes

To assess how homologous recombination might affect patterns of population genetic structure in *Ca.* T. endoseptemdiera in comparison to co-occurring symbionts of *Alviniconcha* and *Ifremeria* snails, we used three commonly applied methods: CLONALFRAMEML [53], MCORR[54] and RHOMETA [55]. CLONALFRAMEML uses a maximum likelihood approach to calculate recombination rates from MAGs, while the other two programs determine recombination rates directly from metagenomic reads, thereby alleviating potential impacts of strain heterogeneity in MAG-based recombination inference.

For the CLONALFRAMEML analyses we followed the approach by Osvatic et al. [56]. *Alviniconcha* and *Ifremeria* symbiont MAGs for this analysis were retrieved from Breusing et al. [7]. We chose the eight most complete MAGs across the geographic range of each symbiont species as this represented the lowest number of genomes available for a single symbiont. MAGs of each symbiont species were first aligned in PROGRESSIVEMAUVE [57] with default parameters. Core locally colinear blocks (LCBs) ≧ 500 bp were then extracted with *stripSubsetLCBs* [57], concatenated, and realigned with MAFFT [58] in auto mode. The core LCB alignment was trimmed with TRIMAL [59] (parameters: -resoverlap 0.75 -seqoverlap 0.80) before phylogenetic reconstruction in RAxML [60] (parameters: -f a -m GTRGAMMA -p 12345 -x 12345 -# 100). The best RAxML tree and the trimmed core LCB alignment were subsequently used in CLONALFRAMEML to infer homologous recombination events among symbiont MAGs. All CLONALFRAMEML analyses were run with 100 pseudo-bootstrap replicates and applying the transition/transversion rate ratios from RAxML as estimates for the kappa parameter. To evaluate the algorithm’s performance, we calculated the compound parameter Rδ [53]. Rδ values greater than 1 typically imply that the relative rate of recombination to mutation R/θ is underestimated due to repeated recombination events at any given genomic position that are not accounted for by the program [53]. R values for the calculation of Rδ were inferred by multiplying R/θ with an estimate of Watterson’s θ according to the following formula: θ = L×G/a_n_, where L is the sum of branch lengths in the recombination-corrected phylogeny, G is the alignment length, and a_n_ is the (n-1)th harmonic number with n being the number of MAGs.

For calculations of recombination and mutation rates from metagenomic reads with MCORR and RHOMETA we used the core and pangenomes of each symbiont species as reference for mapping, respectively. Metagenomic datasets and symbiont pangenomes for *Alviniconcha* and *Ifremeria* for this analysis were retrieved from ref. [7]. For each symbiont species, analyses were run both across and within vent sites to get global and local estimates of recombination to mutation rate ratios. Data were normalized to the same genome coverage across species and the same number of samples per vent site to avoid bias in representation of symbiont strains. For all local analyses, four samples per vent site were used, except in the case of ABE for the *Alviniconcha* Gamma1 symbiont where only three samples were available. For all global analyses, a total of 12 samples were used. All MCORR analyses were run with default settings. However, as the resulting Ф_pool_ and γ/μ values appeared to be unrealistically high, likely due to a poor fit of MCORR’s coalescent model to our data, these results were not used for further evaluation. For the RHOMETA analyses, the mean θ values estimated by the theta_est.nf pipeline were used as input for the creation of lookup tables and inference of the recombination rate parameter ρ. Lookup tables were generated for read depths of 3 to 85 for rho values from 1 to 100 by default. Rho values were divided by the recombination tract length (1000 bp) to get an estimate of recombination rate per site. Recombination to mutation effect ratios were determined according to Krishnan et al. [55] by multiplying the per site ρ/θ ratios with the recombination tract lengths and substitution probabilities. Substitution probabilities were inferred by dividing the number of filtered SNP sites by the pangenome length of each symbiont species.

## Results

### Host populations are undifferentiated across geographic localities

Co-assembly of representative mussel gill samples from Tu’i Malila, ABE and Tahi Moana resulted in 35,060 transcripts that provided a reference for SNP estimation among host populations (Table S3). The curated *B. septemdierum* gill transcriptome assembly comprised an overall size of 57.71 Mb and was 88.90% complete. Assessment of variant sites based on mappings of an average of 50,253,507 cleaned metagenomic reads per sample against the reference transcriptome resulted in 279 SNPs for analysis of host population genomic subdivision. Lowering the minimum sample calling threshold from 75% to 66% recovered 5,639 SNPs that produced equivalent results in terms of population structure. Both principal component analyses based on host covariance matrices and pairwise F_ST_s between vent localities (≦0.0139) indicated an absence of geographic structuring in *B. septemdierum* and a more individual-based partitioning of genomic diversity (Fig. 1B; Table 1). These patterns were confirmed by Mantel tests that indicated no correlation between host identity-by-state and geographic distance matrices (Table 2).

**Table 1.** F_ST_ values between host (above diagonal) and symbiont (below diagonal) populations from different vent sites based on genomic variants. For the symbiont, the pangenome fixation index (P_ST_) based on gene diversities is given in brackets.

|  | Tow Cam | Tahi Moana | ABE | Tu'i Malila |
| --- | --- | --- | --- | --- |
| Tow Cam | * | 0.0139 | 0.0049 | 0.0063 |
| Tahi Moana | 0.1818 (0.1172) | * | -0.0004 | 0.0021 |
| ABE | 0.1888 (0.0753) | 0.0677 (0.1012) | * | -0.0002 |
| Tu'i Malila | 0.2949 (0.1250) | 0.1896 (0.1465) | 0.1678 (0.0614) | * |

**Table 2.**
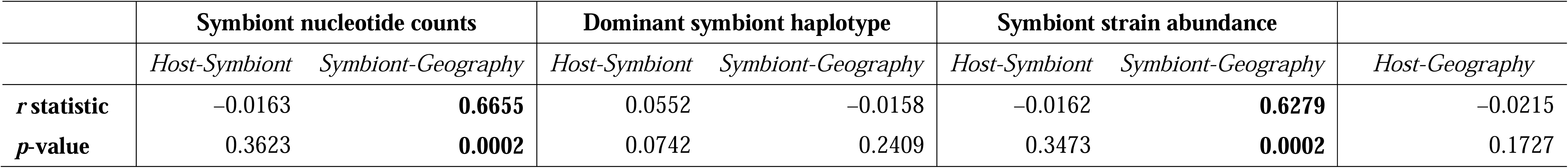
Mantel tests between host, symbiont and geographic distance matrices. Significant tests are indicated in bold.

|  | Symbiont nucleotide counts |  | Dominant symbiont haplotype |  | Symbiont strain abundance |  |  |
| --- | --- | --- | --- | --- | --- | --- | --- |
|  | Host-Symbiont | Symbiont-Geography | Host-Symbiont | Symbiont-Geography | Host-Symbiont | Symbiont-Geography | Host-Geography |
| *r* statistic | -0.0163 | **0.6655** | 0.0552 | -0.0158 | -0.0162 | **0.6279** | -0.0215 |
| *p*-value | 0.3623 | **0.0002** | 0.0742 | 0.2409 | 0.3473 | **0.0002** | 0.1727 |

### Symbiont populations show genetic structure across vent locations

Ninety-one species-level (ANI > 95%), moderate-to high-quality MAGs were combined to reconstruct a ∼3.26 Mb pangenome for *Ca.* T. endoseptemdiera, which consisted of 1040 core and 4352 accessory genes (Table S4, S5). The MAGs underlying this pangenome ranged from 1.75 to 3.50 Mbp in size, had 374–2257 contigs and N50 values of 1836–9202 bp (Table S4). Read mapping against the pangenome reference yielded 2,254 variant sites (SNPs and indels) for symbiont population genomic analyses. Principal coordinate analyses and F_ST_ calculations based on these variant sites indicated moderate partitioning of symbiont populations among vent localities, while P_ST_ indices based on gene diversities suggested weaker differentiation (Fig. 1C; Table 1: F_ST_ = 0.0677–0.2949, P_ST_ = 0.0614–0.1465). Maximum divergence based on pairwise F_ST_s was observed between Tow Cam and Tu’i Malila in accordance with the large geographic distance between these sites, whereas P_ST_ indices did not show an obvious association with geography (Table 1). Overall the degree of population differentiation in *Ca.* Thiodubiliella appeared to be similar to that of the *Ifremeria* SOX symbiont (F_ST_ = 0.0359– 0.3034), but was notably lower than in the co-occurring *Alviniconcha* Epsilon (F_ST_ = 0.1012– 0.8682) and Gamma1 (F_ST_ = 0.2892– 0.5589) symbionts within the ELSC [7].

Intra-host nucleotide diversities were comparable among vent sites, except in the case of Tow Cam which showed significantly lower π values than Tu’i Malila and ABE (Pairwise t-test: p < 0.000002; Fig. S1), resulting in a higher distinctiveness of this site compared to others. Similar to patterns in the sympatric *Ifremeria* and *Alviniconcha* symbionts, the association of *Ca.* Thiodubiliella nucleotide and strain composition with vent locality was significant based on traditional and partial Mantel tests (Table 2; p ≦ 0.002; r ≧ 0.6279), while no correlation between symbiont and host genetic distance could be found. In contrast to the vent snail symbioses, however, the dominant symbiont haplotype appeared to be unrelated to geography (Table 2). Compared to levels of geographic subdivision, inter-sample symbiont population differentiation was notably higher (Fig. 1D; Table S6: F_ST_ = 0.1708–0.7323, P_ST_ = 0.0231– 0.4035), suggesting that symbiont genetic diversity was predominantly partitioned among individual hosts.

### Some symbiont strains are associated with habitat geochemistry or geography

To get more detailed insights into strain composition within and among individual mussel hosts, we decomposed strain haplotypes from the metagenomic dataset with BAYESPATHS. The underlying *Ca*. Thiodubiliella MAG that was binned from the co-assembly and from which strains were decomposed had a total coverage of 12929.84 (40.3–204.4 depth per sample). Fourteen different strains (ANI: 98.88–99.70%) were resolved across samples via this method and the settings chosen, with all strains being detectable in individual hosts – though with drastically varying coverages (Fig. 2; Table S7). Although we cannot fully exclude that the presence of some lowly abundant strains in individual samples might be due to sample cross-contamination during processing or sequencing, analyses of mitochondrial heterogeneity suggested that intra-sample mitochondrial diversity was in general low (0.00–1.32 variants/Kbp) and relatively characteristic of vertically transmitted symbionts and organelles [61, 62]. One mussel individual from ABE exhibited slightly elevated mitochondrial variant densities (3.54 variants/Kbp). The diversity in this sample might reflect natural variability, such as intra-host mutations of the mitochondrial population, possibly due to increased oxidative stress in gill tissue, or – in the case of doubly uniparental inheritance – potential presence of lowly abundant M genomes that leaked into somatic tissue during transmission [50]. Overall, these results indicate that potential sample cross-contamination is likely not an important factor influencing the observed patterns (Table S8). Comparative strain quantifications based on DESMAN were lower than those by BAYESPATHS, recovering only six strains (Fig. S2). Given that the detection of symbiont strains tends to increase with both read depth and number of samples [45], the amount of recovered strains presented here likely provides a snapshot of the true strain diversity within *B. septemdierum*. In addition, due to the high amount of recombination within *Ca.* Thiodubiliella (see below), the detected strains should be regarded as a continuum of possible haplotypes rather than discrete ones, as also confirmed by the distribution of allele frequencies within symbiont populations (Fig. S3). Of the 14 strains identified by BAYESPATHS, one was dominant in almost all individuals regardless of vent site, while others showed a more distinctive distribution according to geochemical habitat and correlated factors (r^2^_adj_ = 0.4617; p = 0.001; Fig. 2A, B). For example, based on redundancy analyses and assessments of intra-host strain abundances, strains 4 and 13 were often relatively frequent in host individuals from Tow Cam, whereas strains 2, 5 and 9 were typically abundant in host individuals from Tu’i Malila (Fig. 2). The association of different strains with geographic location seemed not to be directly linked to their phylogenetic relationships as more closely related strains were not necessarily abundant in the same habitats (Fig. 2B). Given the dominance of a single symbiont strain at all vent sites, no evidence for local adaptation could be found, as correlations between consensus variants (representative of the most abundant strain) and environment were insignificant.

**Fig. 2.**
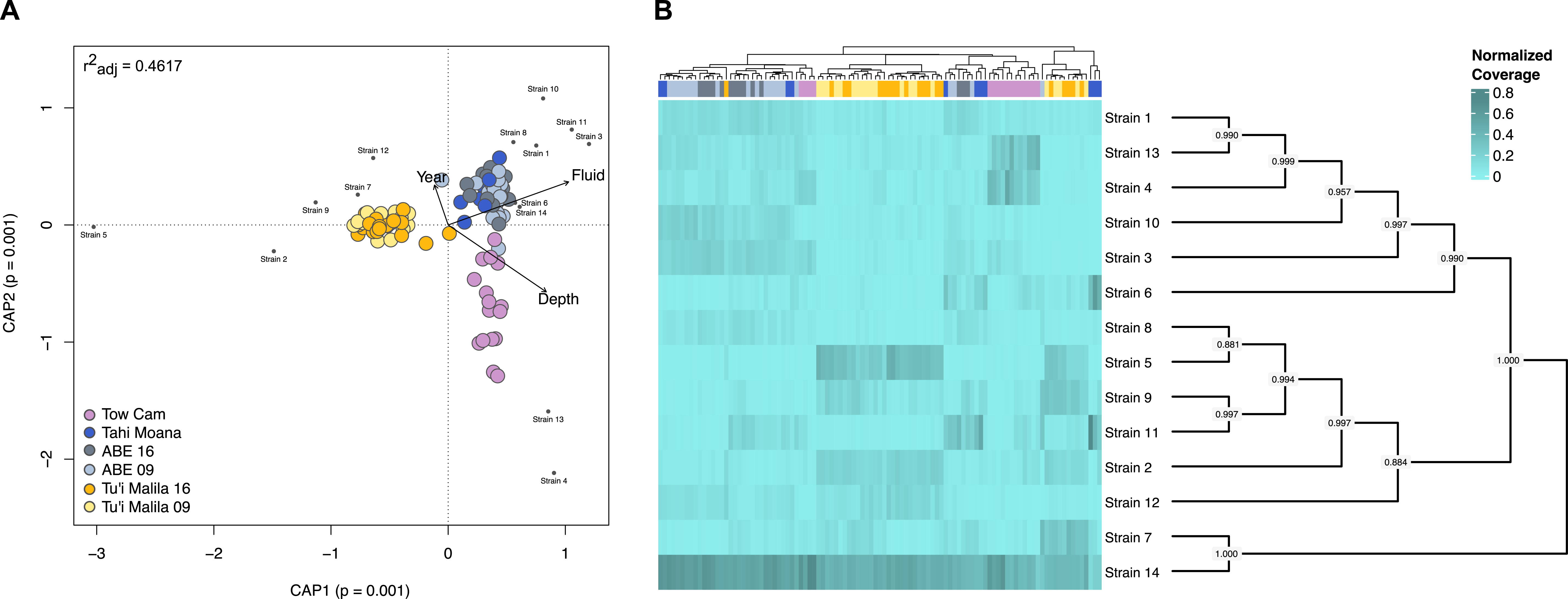
Environmental associations (A) and intra-host abundances (B) of decomposed *Ca.* Thiodubiliella endoseptemdiera strains. Although one symbiont strain is clearly dominant irrespective of sampling locality, some strains are associated with fluid geochemistry or geography of the different vent sites, with strains 4 and 13 being relatively frequent in host individuals from Tow Cam, and strains 2, 5 and 9 being abundant in host individuals from Tu’i Malila. The environmental associations among the distinct symbiont strains are not necessarily related to phylogenetic proximity. Phylogenetic relationships are depicted as a mid-point rooted cladogram. Node values show local bootstrap support.

### Symbiont populations from different vent sites exhibit variation in gene content

We profiled the metagenomic samples using pangenome-based phylogenomic analyses to determine potential gene content variation among symbiont populations from distinct vent sites. At a gene presence/absence threshold of ∼90% between symbiont populations, the main differences were observed for genes with unknown function, followed by genes involved in DNA metabolism and antiviral defense. A small subset of differentially conserved genes was associated with other metabolic categories (Fig. 3; Table S9). For example, *Ca.* T. endoseptemdiera strains from Tow Cam, Tahi Moana and ABE contained a couple of hydrogenase maturation and assembly factors and a heme synthase that were rare in populations from Tu’i Malila (Table S9). By contrast, strains from Tu’i Malila typically encoded a filamentous hemagglutinin, a copper metabolism related gene and an RNA polymerase that decreased in frequency at ABE and Tahi Moana and were almost absent at Tow Cam (Table S9).

**Fig. 3.**
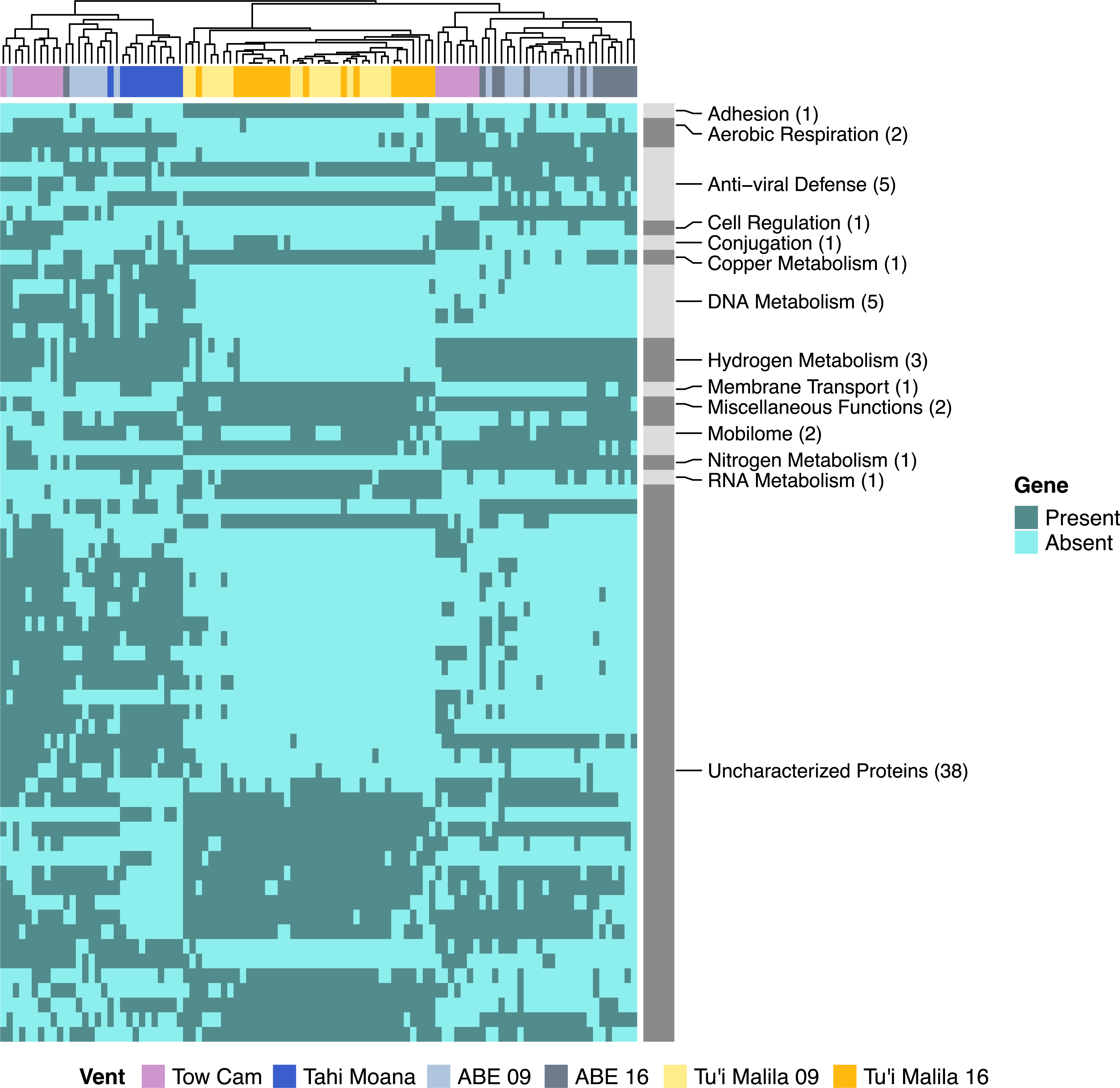
Gene content differences among *Ca.* Thiodubiliella endoseptemdiera populations from vent locations in the Eastern Lau Spreading Center. Samples are ordered based on similarity. Symbiont populations appear to differ mostly in genes associated with unknown functions, DNA metabolism and anti-viral defense, with smaller variation being observed in other metabolic categories. Overall, gene content is more similar among the northern populations at Tow Cam, Tahi Moana and ABE compared to the southernmost population at Tu’i Malila.

### Homologous recombination contributes to cohesiveness of Ca. T. endoseptemdiera populations

Intra-specific homologous recombination plays a critical role in maintaining genetic connectivity of bacterial populations [63]. To evaluate if homologous recombination might have a homogenizing effect on *Ca.* T. endoseptemdiera population structure compared to co-occurring *Alviniconcha* and *Ifremeria* symbiont species [7], we calculated the relative rates and importance of recombination to mutation events from core genome alignments and metagenomic read datasets of each symbiont group. R/θ ratios in core LCB alignments (1,111,427–2,480,742 bp) as assessed by CLONALFRAMEML were at least four times higher in *Ca.* T. endoseptemdiera (0.44) than in thiotrophic symbionts associated with *Alviniconcha* (0.04–0.10) and *Ifremeria* (0.10), while r/m ratios were at least 2.5 times larger (2.38 compared to 0.28–0.94) (Table 3), suggesting pervasive genome-wide recombination within *Ca*. Thiodubiliella (Fig. S4). Although all Rδ values were markedly larger than 1 (Table 3) and CLONALFRAMEML is therefore likely to underestimate R/θ within the inferred parameter ranges [53], the relative ratios among symbiont species should be comparable. Equivalent inferences of recombination parameters from metagenomic reads with RHOMETA suggested relatively high recombination to mutation rate and effect ratios in all analyzed symbionts (Table 3; Fig. 4). Global ρ/θ and r/m values were largest in the Gamma1 symbiont, followed by the two other gammaproteobacterial symbionts (*Ca.* Thiodubiliella and Ifr-SOX) (Table 3; Fig. 4). Local estimates of ρ/θ and r/m showed notable variation between sites for each symbiont species. Median per-site ρ/θ values were slightly increased in the *Ifremeria* SOX symbiont compared to *Ca.* Thiodubiliella and the *Alviniconcha* symbionts, while local recombination effect ratios showed the inverse pattern (Table 3, S10; Fig. 4). Similar to patterns of intra-host nucleotide diversities, the Tow Cam population of *Ca*. Thiodubiliella had lower ρ/θ and r/m values than the other populations, further supporting a higher genetic distinctiveness of this population (Table S10).

**Fig. 4.**
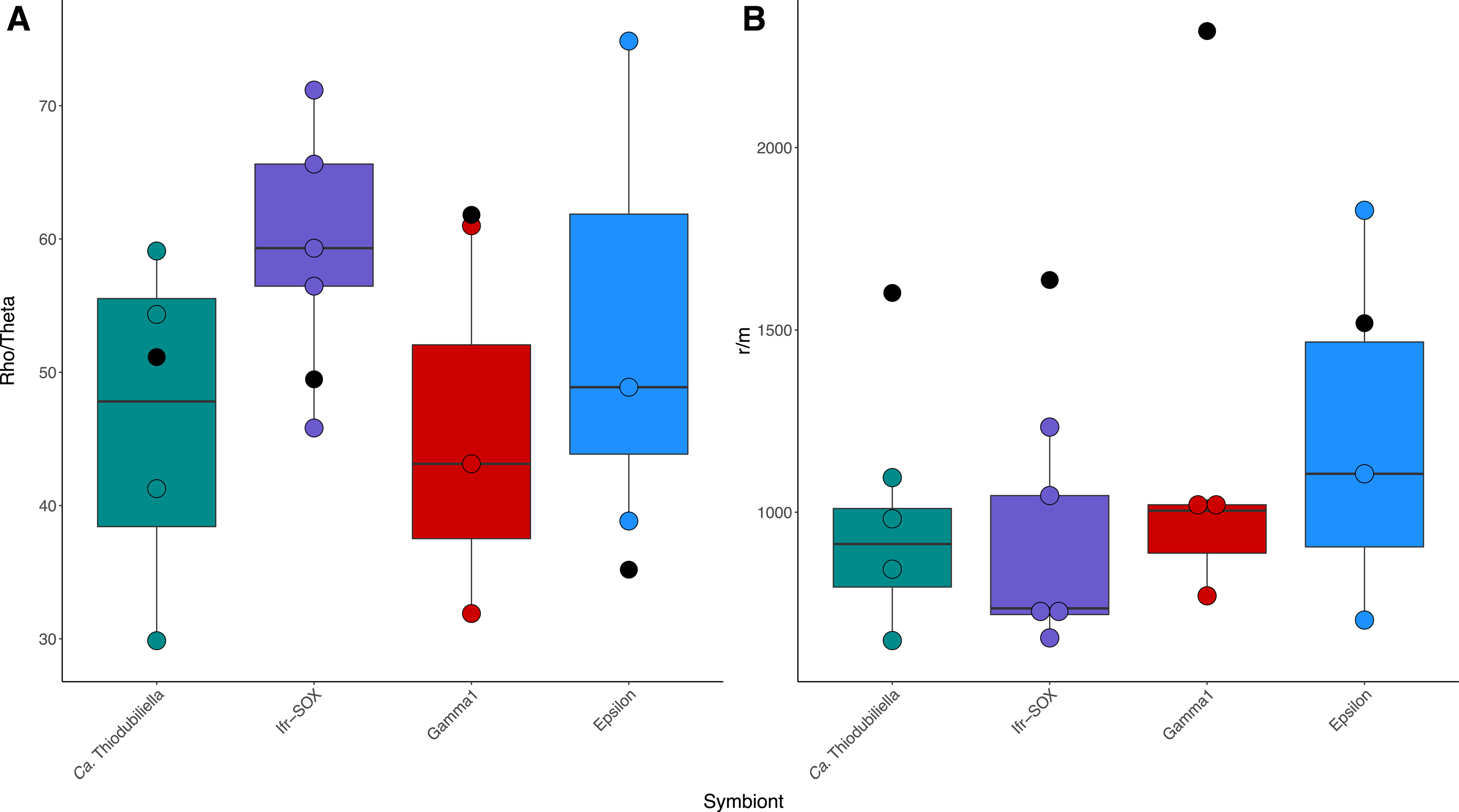
Local (within vent site) recombination rate (A) and effect (B) ratios for chemosynthetic symbionts of *Bathymodiolus*, *Alviniconcha* and *Ifremeria* species from the Lau Basin. Global estimates (across vent sites) are superimposed on the boxplots as black dots.

**Table 3.** Rate and effect ratios of homologous recombination to mutation events in dominant chemosynthetic symbionts of *Bathymodiolus septemdierum*, *Alviniconcha* spp. and *Ifremeria nautilei* from the Lau Basin and Tonga Volcanic Arc. Estimates are given for inferences with i) ClonalFrameML based on MAGs and ii) Rhometa based on metagenomic read alignments. For the ClonalFrameML analyses a 95% confidence interval is given for each ratio and alignment lengths for each symbiont are reported, with the percentage of the pangenome lengths provided in brackets. For the Rhometa analyses, both global (across sites within species) and local (within sites within species) estimates of recombination are shown. R/θ, ρ/θ = recombination to mutation rate ratio inferred by ClonalFrameML and Rhometa, respectively. r/m = relative effects of recombination to mutation. Rδ = key determinant for algorithm performance in ClonalFrameML.

|  | *Ca. Thiodubiliella* (*B. septemdierum*) | *Epsilon* (*A. boucheti*) | *Gamma1* (*A. kojimai*, *A. strummeri*) | *Ifr-SOX* (*I. nautili*) |
| --- | --- | --- | --- | --- |
| CLONALFRAMEML |  |  |  |  |
| # MAGs | 8 | 8 | 8 | 8 |
| Alignment length [bp] | 1,111,427 (34.11%) | 1,152,639 (65.32%) | 2,090,501 (51.74%) | 2,480,742 (51.40%) |
| LCBs | 550 | 672 | 895 | 998 |
| R/θ [95% CI] | 0.44 (0.42–0.45) | 0.10 (0.10–0.11) | 0.04 (0.03–0.05) | 0.10 (0.09–0.11) |
| r/m [95% CI] | 2.38 (2.23–2.53) | 0.28 (0.25–0.30) | 0.58 (0.48–0.70) | 0.94 (0.81–1.09) |
| Rδ | 85247.70 | 14109.17 | 7892.98 | 7713.08 |
| **RHOMETA** |  |  |  |  |
| *Global* |  |  |  |  |
| # samples | 12 | 12 | 12 | 12 |
| Pangenome size [bp] | 3,258,339 | 1,764,546 | 4,040,567 | 4,826,737 |
| ρ/θ | 51.14 | 35.19 | 61.81 | 49.47 |
| r/m | 1601.42 | 1518.53 | 2319.64 | 1636.89 |
| *Local* |  |  |  |  |
| # samples per site | 4 | 4 | 3–4 | 4 |
| Pangenome size [bp] | 3,258,339 | 1,764,546 | 4,040,567 | 4,826,737 |
| Median ρ/θ [range] | 47.80 (29.86–59.10) | 48.88 (38.84–74.85) | 43.13 (31.90–60.99) | 59.31 (45.82–71.17) |
| Median r/m [range] | 912.94 (648.19–1095.06) | 1105.59 (704.38–1827.99) | 1004.44 (770.89–1036.48) | 736.52 (655.45–1233.57) |

## Discussion

Symbiont strain composition in horizontally transmitted symbioses is determined by a variety of interacting factors, including host genetics, environment and geography [2]. At deep-sea hydrothermal vent sites, local adaptation to geographically variable fluid geochemistry has been shown to play a significant role in shaping strain-level variation of chemosynthetic endosymbionts [7]. It is currently unknown how these patterns are affected by niche differences among symbiotic associations that could influence the degree of selective pressures on both host and symbiont. To address this question, we assessed symbiont strain composition in populations of the deep-sea mussel *Bathymodiolus septemdierum* and compared our results to previous findings in co-occurring vent snail symbioses [7] that segregate into contrasting ecological niches [19].

Symbiont strain composition in *B. septemdierum* was unrelated to host genetics but varied to some degree with vent habitat and geography. Although there are clear similarities to the population genomic structures of sympatric *Alviniconcha* and *Ifremeria* symbionts given that strain proportions were distinct among vent sites, the dominant *Ca.* Thiodubiliella endoseptemdiera strain was contrastingly shared across localities without any evidence for local adaptation. While we cannot directly determine the cause of the difference among the snail and mussel symbionts given phylogenetic constraints and interlinked ecological variables that differ among these taxa, there are notable habitat and feeding differences between the snails and mussel species that could play a role. For example, the lower thermal and geochemical variation in the microhabitat patches that *B. septemdierum* symbioses occupy [19, 20] might favor one main *Ca.* T. endoseptemdiera strain across the symbiont’s distributional range, thus allowing host individuals to affiliate with the same symbiont genotype at different vent locations. In addition, the increased mixotrophic capacity of *Bathymodiolus* relative to other chemosymbiotic vent invertebrates, including *Alviniconcha* and *Ifremeria*, could reduce dependence on optimal symbiont function and, thus, selection for a locally adapted strain, similar to what has been suggested for facultative host-microbe associations [64]. *Bathymodiolus* species have a functional albeit reduced digestive tract and can complement their symbiont-derived nutrition with particulate and dissolved organic matter from their surroundings [65, 66], which is thought to facilitate segregation into local habitat patches with barely detectable chemosynthetic reductants [19, 20]. In fact, with sufficient food, bathymodioline mussels have been observed to survive without their symbionts for months to possibly years, at least in a laboratory setting [67]. By contrast, there is no evidence that *Alviniconcha* and *Ifremeria* snails rely significantly on heterotrophic feeding, though they still retain reduced and functional digestive systems [68–70].

Despite absence of habitat affiliation for the dominant *Ca.* T. endoseptemdiera strain, symbiont populations exhibited differences in gene content and abundances of minority strains, indicating that environmental conditions might at least partially shape the population genetic structure of *Ca.* T. endoseptemdiera. For example, the strains from Tu’i Malila typically missed several hydrogenase maturation factors that were present in the northern populations, possibly in adaptation to the low hydrogen concentrations at this site. In addition, we observed some gene content differences related to anti-viral defense. In line with the strong host and habitat restriction of viruses at hydrothermal vents [71], the composition of viral communities might be an important and possibly common mediator of symbiont genetic differentiation in the deep sea. Similar impacts have been reported for symbiont populations associated with siboglinid tubeworms [11, 72] and vent-associated snails [7].

We found that up to 14 distinct strains co-occurred within a single mussel individual, although the actual number of strains is likely higher, given that strain detection depends notably on read coverage, number of samples, sequencing method and bioinformatic algorithm assumptions. In general, however, these estimates are within the range of variation reported for thiotrophic symbionts of other *Bathymodiolus* species, including *B. brooksi* from cold seeps in the Gulf of Mexico (up to 9 strains) [39] and *B. puteoserpentis* from hydrothermal vents on the Mid-Atlantic Ridge (up to 16 strains) [73]. Symbiont diversity in bathymodioline mussels has previously been shown to be maintained by functional heterogeneity among symbiont subpopulations and is thought to confer metabolic flexibility of the symbiosis to changing environmental conditions [73, 74]. Our data strongly support this hypothesis, suggesting that *B. septemdierum* hosts retain one, broadly well-performing symbiont strain across their distributional range and complement this strain by local, habitat-specific symbiont genotypes with a variety of functions.

The widespread occurrence of one *Ca.* Thiodubiliella strain across vent localities indicates that dispersal barriers are likely not the main drivers of symbiont population structure, although overall population differentiation tended to increase with geographic distance (or correlated factors). These patterns are in contrast to those in other *Bathymodiolus* symbionts where geography is thought to be a major factor underlying population subdivision [12, 13], at least over much larger spatial scales (1000s of km) than assessed here. It is possible that the comparatively small distance among vent sites in the Lau Basin (<200 km), as well as contrasts in oceanographic conditions or symbiont properties, contribute to these differences. Recent biophysical models for the Lau Basin have implied that dispersal barriers within this region are mostly absent [75], while evidence from a variety of biogeographic studies suggests that vent-associated microbial communities are not significantly shaped by dispersal limitations [76].

However, the dispersal mechanisms of bacterial symbionts are not well understood, making it difficult to assess how well dispersal patterns are represented by model simulations. In addition, several population genetic studies on the symbionts of vent-associated animals, including those of *Bathymodiolus* species from mid-ocean ridge systems and other mollusk taxa from the Lau Basin, have highlighted the effect of topological barriers and geographic distance on symbiont biogeography [7, 12, 13]. Despite contrasting evidence from these systems, it is possible that *Ca*. T. endoseptemdiera faces less dispersal restrictions than co-occurring symbionts by employing different dispersal strategies. For example, the bacterial symbionts of lucinid clams have been hypothesized to be transported over long distances with seagrass fragments [56], whereas some algal symbionts of gorgonian octocorals are known to hitchhike with the corals’ larvae [77] – a strategy that has also been suggested for the symbionts of bathymodioline mussels [78]. Considering the broad distribution of *B. septemdierum* in the Western Pacific and Indian Ocean [16], vectored transport, if it can be confirmed, would greatly increase the dispersal ability of *Ca.* T. endoseptemdiera, thereby reducing geographic partitioning among disparate populations. It is currently unknown how widely distributed *Ca.* T. endoseptemdiera is, as sequence information for bacterial symbionts of other *B. septemdierum* populations is scarce. Based on comparisons of publicly available full-length 16S rRNA sequences, *Ca.* T. endoseptemdiera from the Lau Basin is >97% identical to other *B. septemdierum* symbionts from the Izu-Ogasawara Arc vents in the Northwest Pacific. In addition, our sequence searches within the Integrated Microbial NGS platform (https://www.imngs.org) [79] at an identity cutoff of 100% suggest that *Ca.* T. endoseptemdiera might occur at various locations in the Northeast Pacific Ocean and around the Solomon Islands. Although further genomic analyses will be needed to confirm the species-level designations of the recovered sequences, these results support a possible widespread distribution of *Ca.* T. endoseptemdiera.

Compared to the genetic subdivision across geographic locations, the degree of symbiont population divergence between individual mussels was approximately 2.5 times higher (based on pairwise F_ST_). The detected levels of inter-host symbiont population differentiation agree with the patterns observed in other sulfur-oxidizing *Bathymodiolus* symbionts and were similarly unrelated to host genotype [13, 39], suggesting that variation in symbiont composition between host individuals is driven by neutral evolutionary processes [39]. Although *Bathymodiolus* mussels have been shown to remain competent for symbiont uptake throughout their lifetime [80], Romero Picazo et al. [39] suggested that strong genetic isolation among individual symbiont populations can occur if environmental symbiont acquisition is restricted in later host developmental stages and re-colonization happens increasingly through self-infection. This model of symbiont transmission might explain how divergence of symbiont populations is more strongly maintained on an individual than on a geographic level. Founder events combined with possible priority effects [81] during symbiont infection may increase inter-host genetic differentiation of symbiont populations. By contrast, occasional reshuffling of symbiont strains between host and environment combined with adequate symbiont dispersal may promote homologous recombination and mixing of the free-living symbiont pools, thereby diminishing geographic and ecological isolation. *Candidatus* T. endoseptemdiera has previously been shown to have some of the highest effective recombination rates observed for bacteria [4]. Our data indicate that the relative rates of recombination to mutation in all analyzed mollusk symbionts were elevated and surpassed an estimated threshold beyond which recombination is theoretically sufficient to prevent population differentiation [82]. In *Ca*. T. endoseptemdiera high recombination rates paired with diminished effects of environmental selection or dispersal barriers might contribute to the lower geographic subdivision of this species compared to sympatric *Alviniconcha* and *Ifremeria* symbionts. Similar observations have been made in chemosynthetic lucinid clam symbioses, where increased levels of homologous recombination are hypothesized to maintain genetic cohesiveness within a cosmopolitan symbiont species [56].

## Conclusions

Our data provide evidence that symbiont populations associated with *Bathymodiolus septemdierum* are structured among vent locations in the Lau Basin, but show no signs for habitat affiliation of the dominant symbiont strain, in sharp contrast to thiotrophic symbionts of co-occurring *Alviniconcha* and *Ifremeria* snails. While the lower dependence of *B. septemdierum* on symbiont-derived nutrition might contribute to these patterns, our current data do not allow disentangling these effects from impacts of correlated ecological partitioning. For example, related to its mixotrophic feeding mode, *B. septemdierum* occupies niches with less variable fluid flows, which might promote affiliations with the same symbiont strain across geographically distinct, but ecologically similar habitats. Our findings further suggest that symbiont-related factors, such as high recombination rates combined with the possibility of long-distance dispersal, might decrease population differentiation among vent locations. However, the relative importance of these processes for symbiont geographic structure and adaptation remains to be determined. Future investigations of environmental versus intra-host strain abundances at different vent sites as well as extracellular presence of symbionts on host larvae will help to shed light on this question.

## Acknowledgements

We thank the captains and crews of the R/Vs *Falkor* and *Thomas G. Thompson* as well as the pilots of the ROVs *Ropos* and *Jason* for their invaluable support of the sample collections. The technical staff at Novogene, Co. and Psomagen, Inc. are gratefully acknowledged for preparing and sequencing the Illumina metagenomic and RNA-seq libraries. We further thank the Kingdom of Tonga for providing access to their national waters and Abi Goodman for her help in verifying the host species identities of mussel samples used in this study. This work was funded by the Key Special Project for Introduced Talents Team of Southern Marine Science and Engineering Guangdong Laboratory (Guangzhou) (GML2019ZD0409), the General Research Fund of Hong Kong SAR (16101219, 16101822), the Schmidt Ocean Institute, the U.S. National Science Foundation (grant numbers OCE-1536331, 1819530 and 1736932 to R.A.B. and EPSCoR Cooperative Agreement OIA-#1655221) and the National Institutes of Health (grant numbers 5K99GM135583-02 to S.L.R. and 5R35GM128932-03 to R.C.D.).

## Data Availability Statement

Raw metagenomic and RNA-seq reads are available in the National Center for Biotechnology Information under BioProject numbers PRJNA855930 and PRJNA849542. The *Bathymodiolus septemdierum* transcriptome has been deposited in the Transcriptome Shotgun Assembly database under accession number GJZH00000000. Host mitochondrial genomes and *Ca.* Thiodubiliella endoseptemdiera MAGs have been submitted to GenBank under accession numbers listed in Supplementary Tables S1 and S4, respectively.

