## Supplementary Material for "Ecological differences among hydrothermal vent symbioses may drive contrasting patterns of symbiont population differentiation"

**This PDF file includes:**

Figure S1 to S4

Legends for Tables S1 to S10

**Other supplementary materials for this manuscript include the following:**

Tables S1 to S10

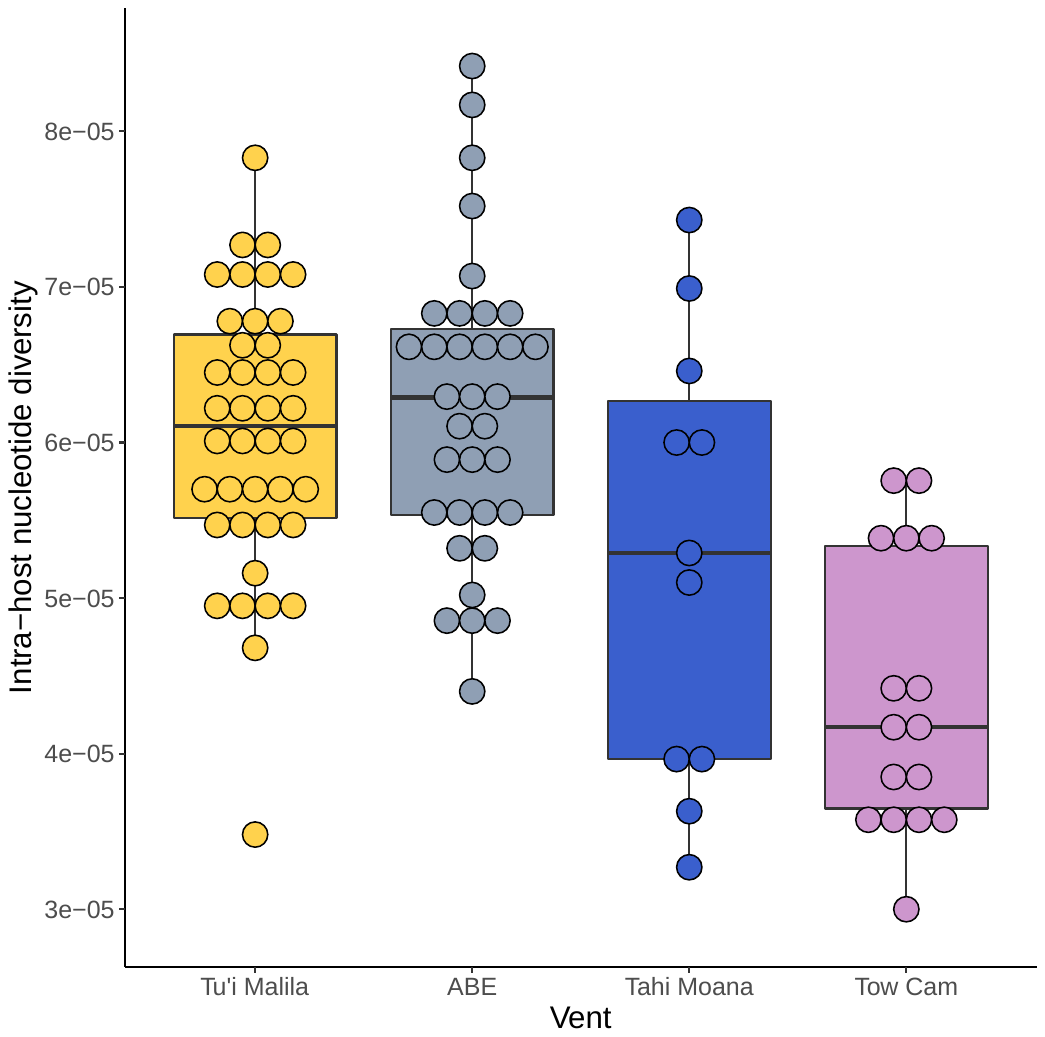

**Fig. S1** Intra-host nucleotide diversities (π) at each vent site. Mean π values are significantly lower at Tow Cam than at ABE and Tu’i Malila based on pairwise t-tests (p < 0.000002).

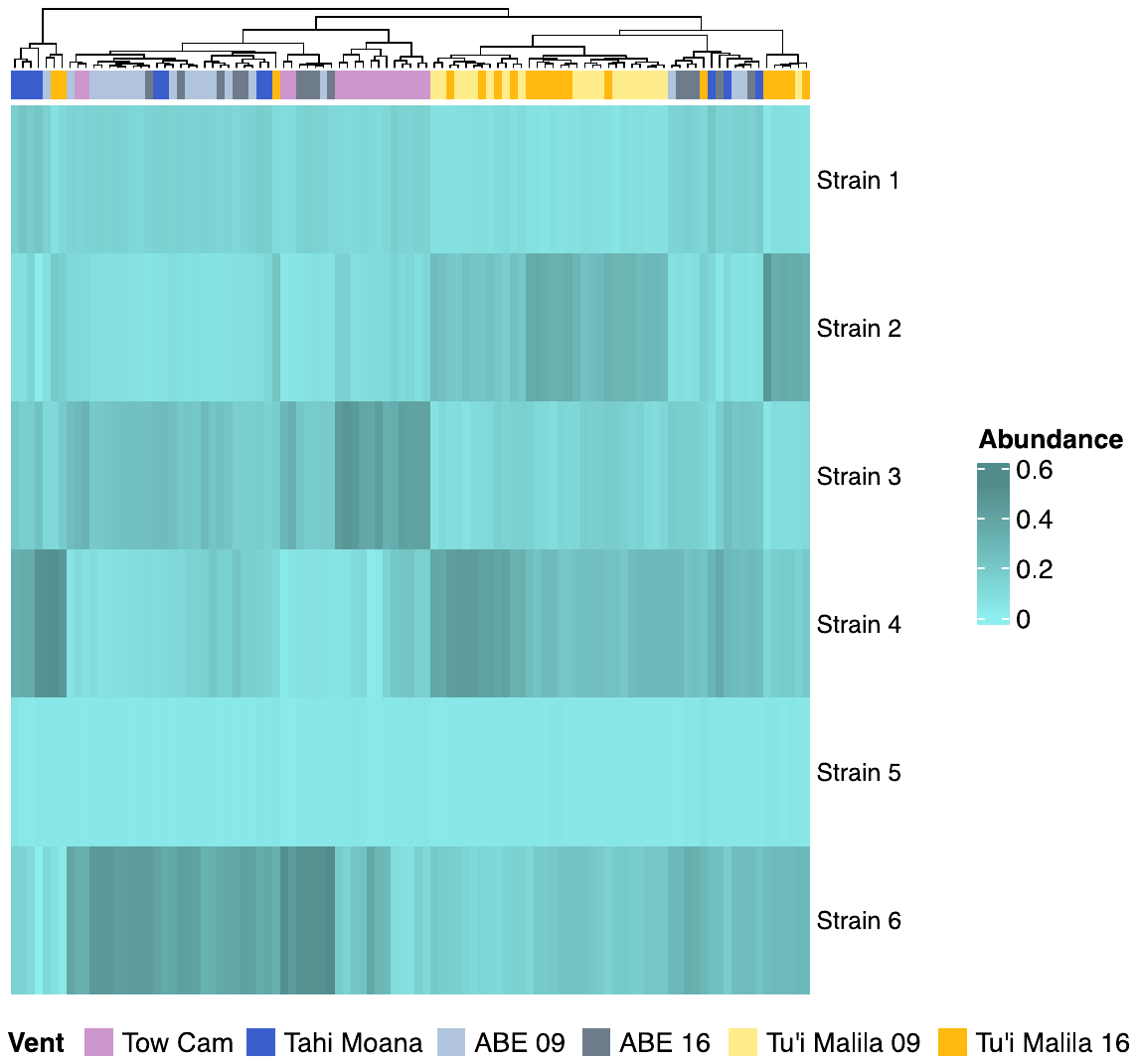

**Fig. S2** Strain composition inferred through haplotype extraction from metagenomic core variant co-occurrence with DESMAN. Although DESMAN detects notably less strains than the BayesPaths algorithm, a similar pattern of strain abundance clustering by vent field, in particular for Tow Cam and Tu’i Malila, is observed with this method.

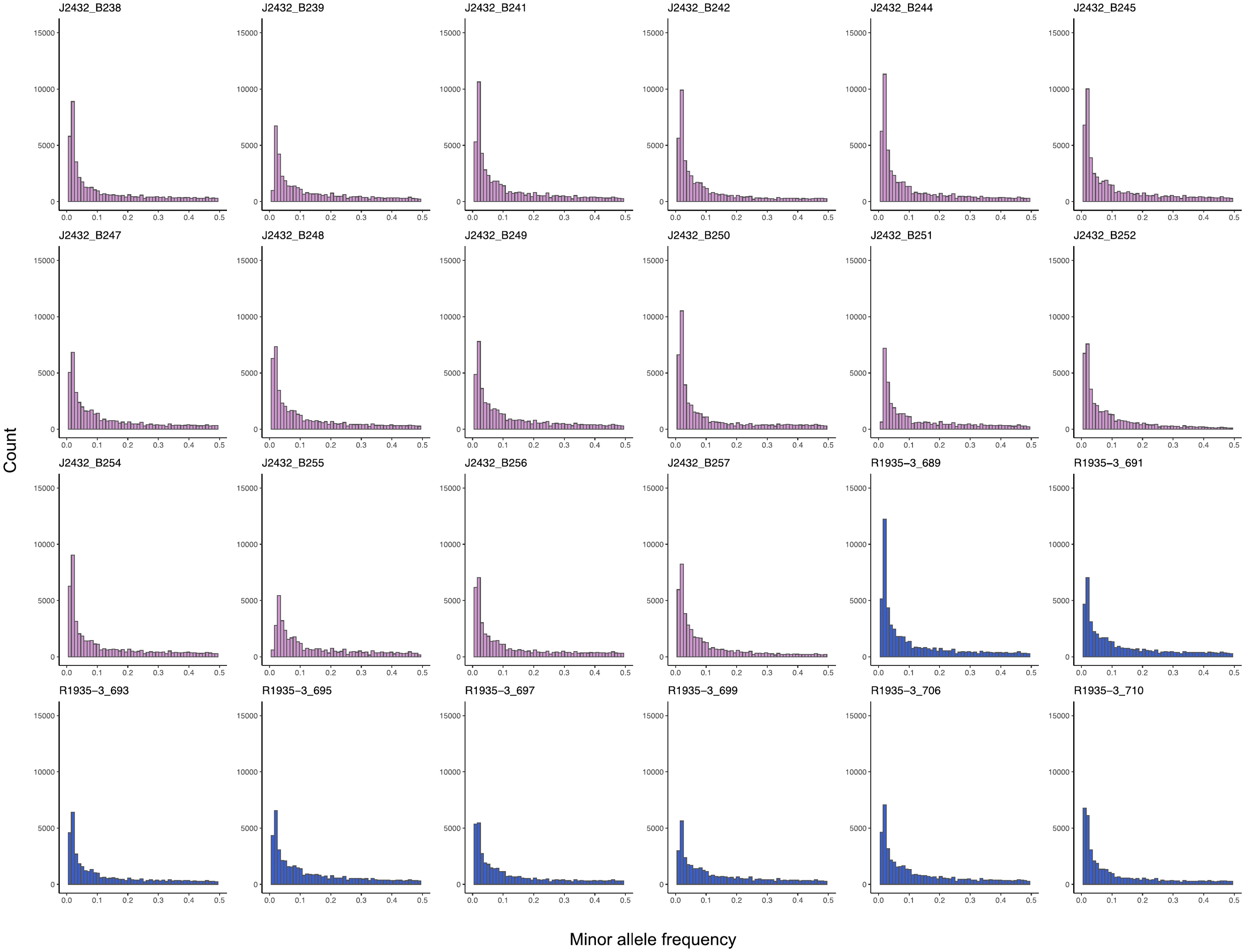

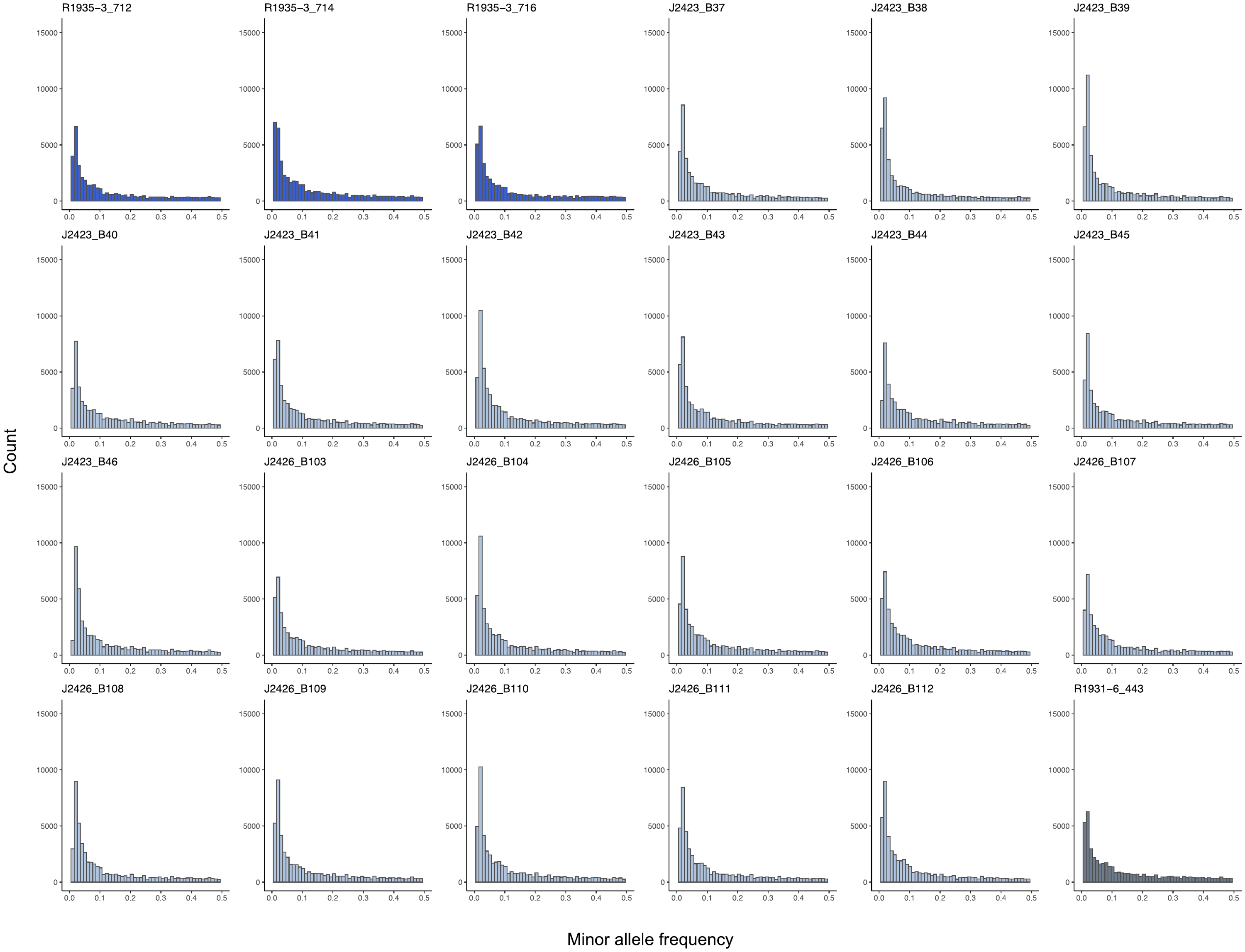

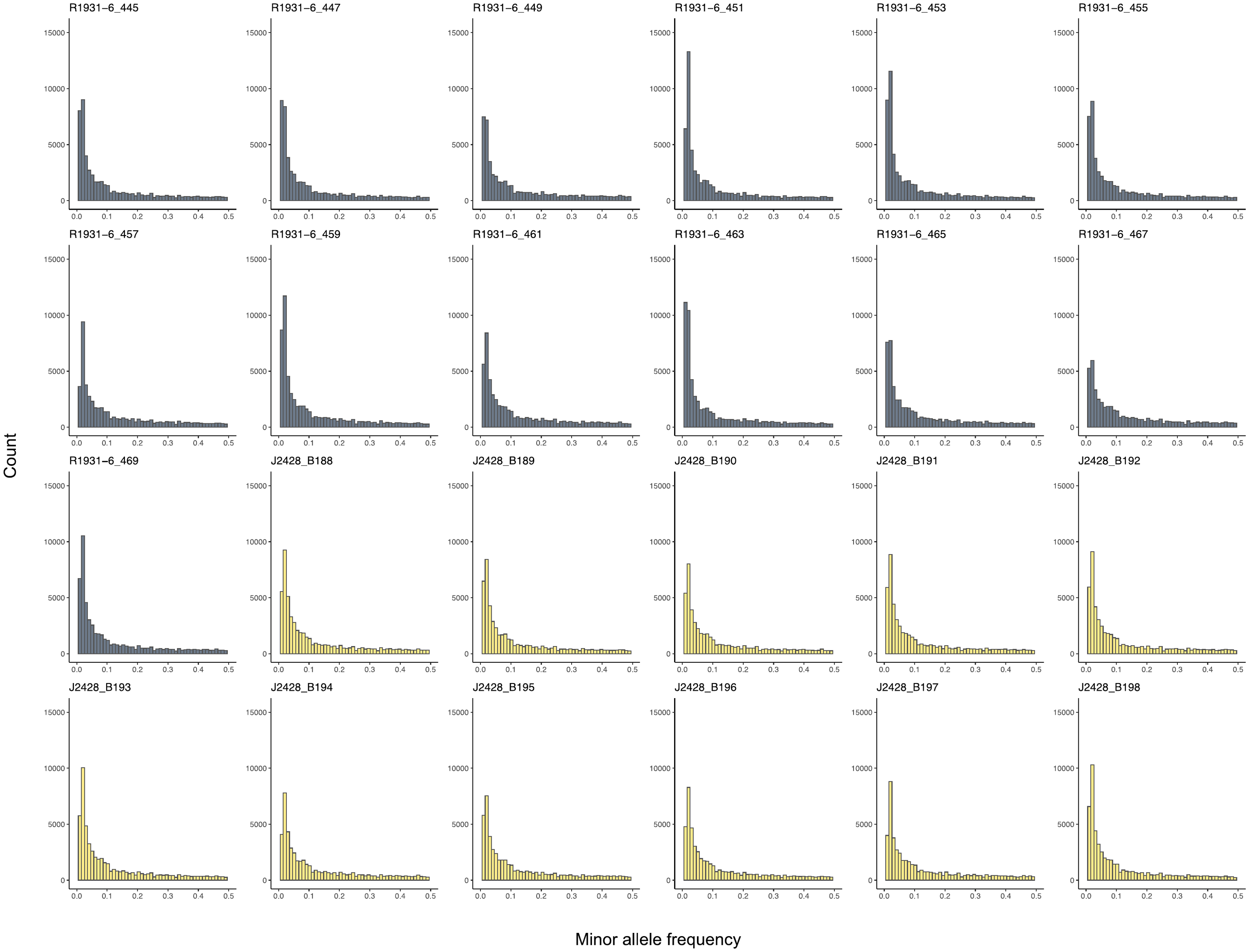

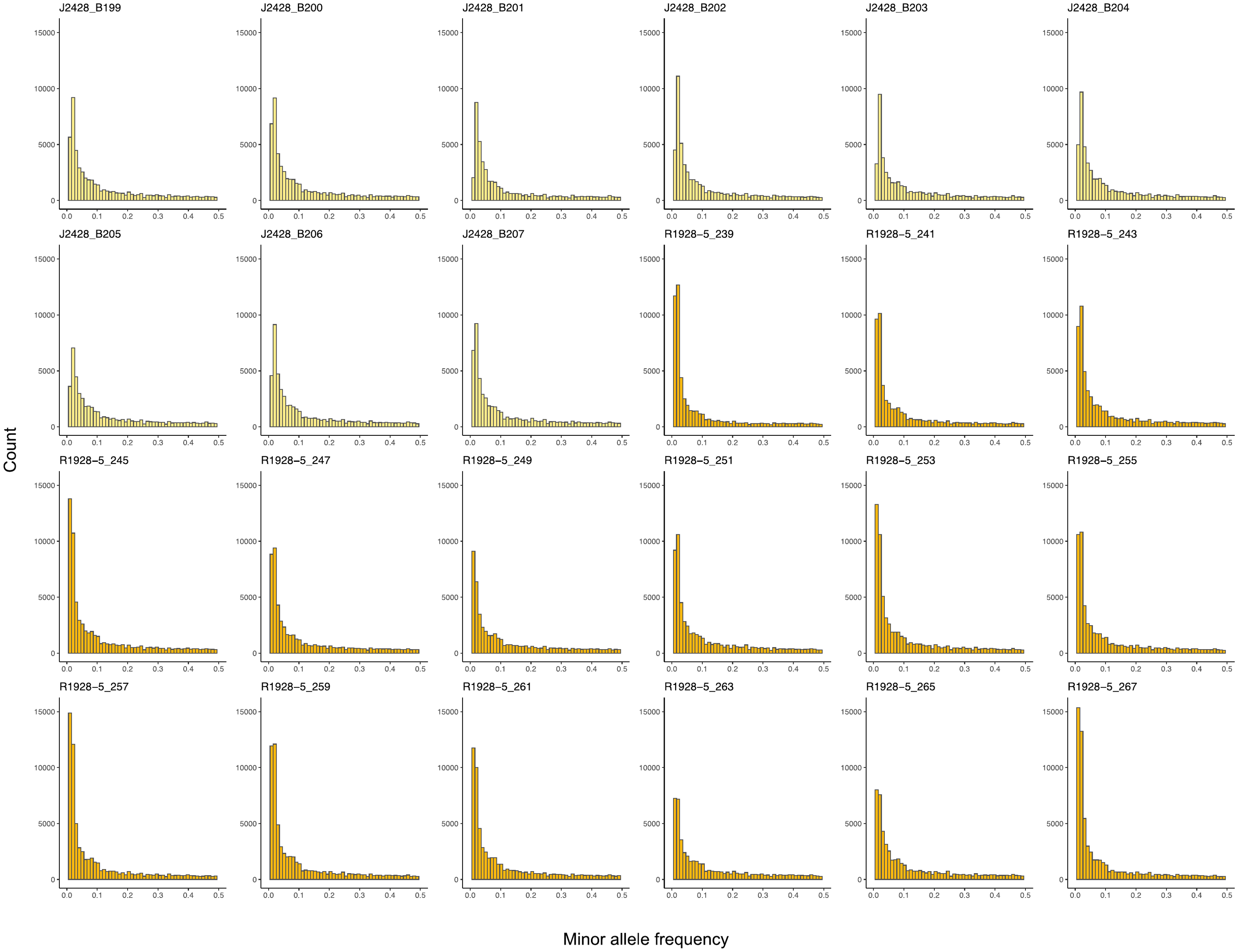

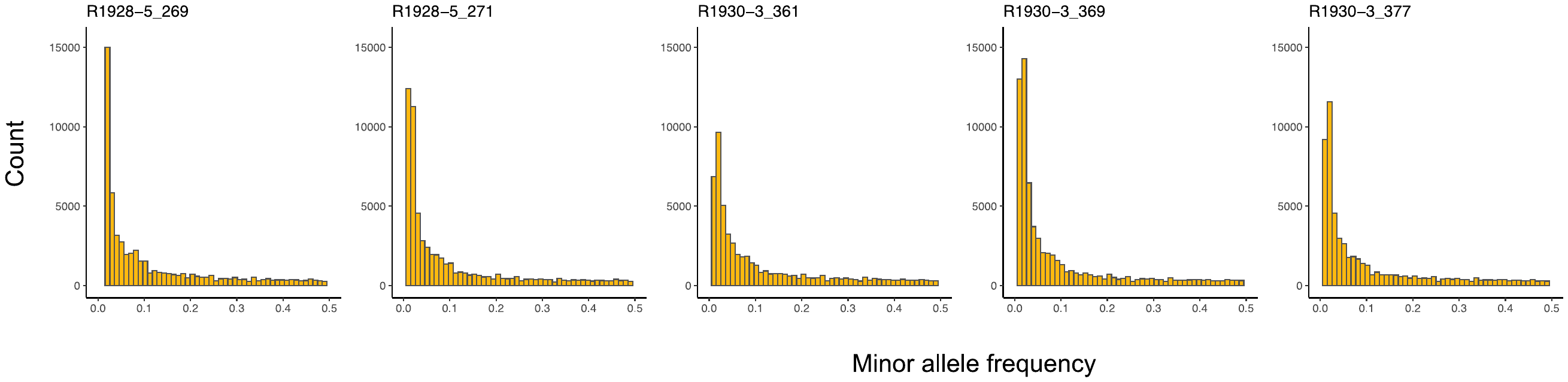

**Fig. S3** Minor allele frequency spectra for each mussel sample. AFSs follow expectations for an equilibrium neutrally-evolving population, in accordance with the high genetic diversity observed for *Ca.* T. endoseptemdiera and other mussel symbionts as previously described [1].

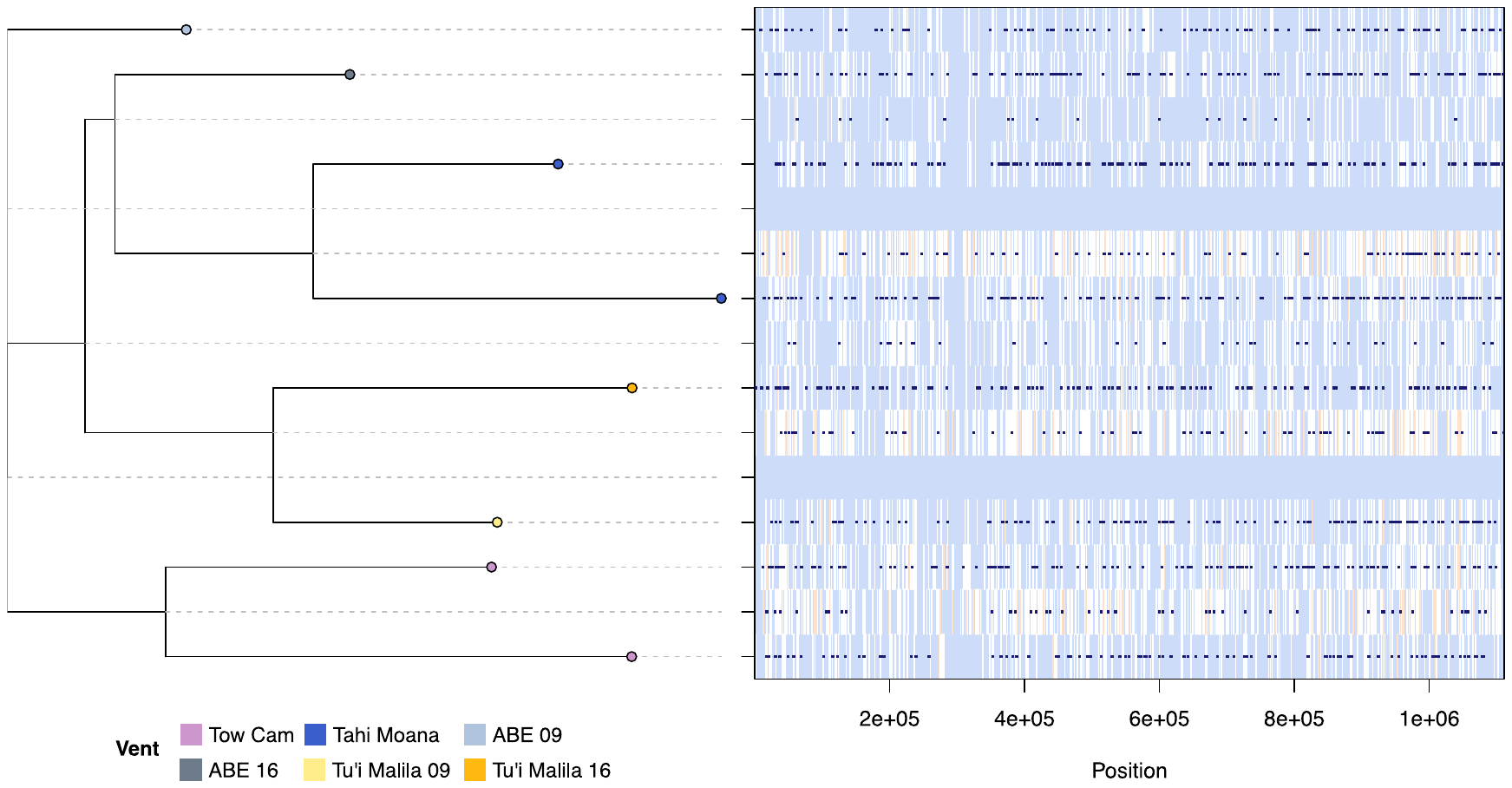

**Fig. S4** Analysis of genomic recombination in *Ca.* Thiodubiliella endoseptemdiera with ClonalFrameML. Left: recombination-corrected phylogeny, with tip colors indicating the isolation source of the symbiont genome. Right: recombination and mutation events across the core genome. Recombination events are shown as midnightblue bars, while mutation events are indicated as white to red bars, with white denoting non-homoplasic substitutions and increasing redness denoting increasing levels of homoplasy. Light purple bars indicate absence of substitutions. Recombination is pervasive across the core genome of *Ca.* Thiodubiliella endoseptemdiera, though rates are likely underestimated due to repeated recombination events at any given genomic position.

**Table S1** Collection information for *Bathymodiolus septemdierum* samples in the Lau Basin.

**Table S2** Effects of methodology and vent habitat on symbiont genetic variation as assessed through PERMANOVAs. PERMANOVAs were performed with the adonis2 function in R.

**Table S3** Overview of the *Bathymodiolus septemdierum* transcriptome assembly.

**Table S4** Quality statistics and taxonomic assignments for metagenome-assembled genomes of *Ca*. Thiodubiliella endoseptemdiera.

**Table S5** Overview of the *Ca*. Thiodubiliella endoseptemdiera pangenome assembly.

**Table S6** Pairwise F_ST_s (lower diagonal) and P_ST_s (upper diagonal) between symbiont populations of individual hosts.

**Table S7** Strain abundances in analyzed mussel hosts. Abundance values represent normalized coverages as reported by STRONG. Non-normalized strain coverages are given in brackets.

**Table S8** Detected mitochondrial variants in each sample. Variants were determined across the whole mitochondrial genome including and excluding the control region, which is typically difficult to assemble and can therefore lead to false variant calls.

**Table S9** Differentially preserved genes between *Ca*. Thiodubiliella endoseptemdiera populations from Tu'i Malila, ABE, Tahi Moana and Tow Cam. For each vent location the proportion of symbiont samples containing the respective gene variant is shown.

**Table S10** Local (vent site-specific) recombination analyses with RHOMETA within each symbiont species. ρ/θ = recombination to mutation rate ratio, r/m = relative effects of recombination to mutation.
